# *Rho* enhancers play unexpectedly minor roles in *Rhodopsin* transcription and rod cell integrity

**DOI:** 10.1101/2022.12.02.518920

**Authors:** Chi Sun, Philip A. Ruzycki, Shiming Chen

## Abstract

Enhancers function with a basal promoter to control the transcription of target genes. Enhancer regulatory activity is often studied using reporter-based transgene assays. However, unmatched results have been reported when selected enhancers are silenced *in situ*. In this study, using genomic deletion analysis in mice, we investigated the roles of two previously identified enhancers and the promoter of the *Rho* gene that codes for the visual pigment rhodopsin. The *Rho* gene is robustly expressed by rod photoreceptors of the retina, and essential for the subcellular structure and visual function of rod photoreceptors. Mutations in *RHO* cause severe vision loss in humans. We found that each *Rho* regulatory region can independently mediate local epigenomic changes, but only the promoter is absolutely required for establishing active *Rho* chromatin configuration and transcription and maintaining the cell integrity and function of rod photoreceptors. To our surprise, two *Rho* enhancers that enable strong promoter activation in reporter assays are largely dispensable for *Rho* expression *in vivo*. Only small and age-dependent impact is detectable when both enhancers are deleted. Our results demonstrate context-dependent roles of enhancers and highlight the importance of studying functions of *cis*-*regulatory regions* in the native genomic context.

## INTRODUCTION

Transcriptional regulation is a tenet of cellular development and homeostasis. Each cell type uses a unique set of regulatory sequences, namely promoters and enhancers, to enact expression of specific genes. A gene’s promoter is the DNA region immediately upstream of the transcription start site (TSS). Enhancers are often found distally upstream or downstream of a gene’s coding region and modulate the promoter activity. The cooperative regulation by promoter and enhancers is important for directing gene transcription within a specific cell type(s) at a precise timing and an appropriate level. In particular, promoters and enhancers contain accessible regions of sequence-specific binding sites for cell-type specific and general transcription factors (TFs). Base-content analysis indicates the differences between the two classes: promoters display high-GC content and contain the requisite TATA box, initiator sequences and binding sites for a variety of TFs; enhancers are often AT-rich and contain binding sites for cell-type specific TFs^1-4^. Both classes can be defined by the presence of active histone modifications; specifically, promoters with H4K4me3 and H3K27Ac marks^5,6^, and enhancers with H3K27Ac and H3K4me1 marks^7^. Studies have further classified the regulatory units based on the landscape and density of the enrichment of these histone modifications. Super enhancers and broad H3K4me3 domains^8-13^ are hence defined as genomic regions regulating the specific gene expression that is imperative to a cell’s state and function.

Rod and cone photoreceptors are the cell types of the retina, and mediate the conversion of light photons into electrical signals in a process called phototransduction. In mouse and human retinas, rod photoreceptors reside within the outer nuclear layer (ONL) and comprise 70% of all retinal cells^14-16^.

Rod photoreceptors make synaptic connections at the outer plexiform layer (OPL) with horizontal neurons and bipolar cells that reside in the inner nuclear layer (INL) to transfer light-evoked signals further to the brain via ganglion cell axons that comprise the optic nerve. The first step of phototransduction is accomplished within a unique cellular structure called the outer segment (OS) which consists of a stack of membrane discs packed with the light-sensing visual pigment, Rhodopsin^17-19^.

Rhodopsin is a G-protein-coupled receptor consisting of an opsin apo-protein and its chromophore 11-*cis*-retinal that accounts for the absorption properties. Rhodopsin is also required for OS structural integrity^20,21^. Mutations in the human *Rhodopsin* gene (*RHO*) cause blinding diseases, such as retinitis pigmentosa (RP) and night blindness ^22-25^. The *Rho* gene is tightly regulated and highly transcribed, with its mRNA representing the most abundant transcript in rod photoreceptors ^26-28^. Irregular *Rho* expression, either at lower or higher than normal levels, can lead to photoreceptor degeneration in animal models ^29-31^. Mechanisms of the precise regulation on *Rho* expression have been extensively investigated. Initial studies identified the *Rho proximal promoter region* (*PPR)*, a conserved ∼200bp region immediately upstream of *Rho* TSS, which was sufficient to drive the expression of the *LacZ* reporter in mouse photoreceptors^32,33^. Addition of a conserved genomic region, the *Rho enhancer region* (*RER*) lying 2kb upstream of *PPR*, greatly increased *PPR*-mediated *LacZ* expression ^32,33^.

Individual *cis*-*regulatory elements (CREs)* within *PPR* are detailed as sequence motifs bound by photoreceptor-specific TFs, including Cone-Rod Homeobox (CRX) ^34,35^ and Neuroretina Leucine Zipper (NRL)^36-38^. ChIP-seq targetome analysis has discovered that CRX and NRL cooperatively bind not only to *PPR* and *RER*, but also to a region 4 kb upstream of *Rho* TSS, which is designated as *CRX-Bound Region 1* (*CBR*)^39,40^. Both *CBR* and *RER* could further activate a *PPR-*driven reporter by 8-10 fold in retinal explant reporter assays^39^. CRX/NRL synergistically activated *PPR* activity in transient HEK293 cell transfection assays, and loss of CRX or NRL in mice caused reduction in *Rho* expression and defects to rod photoreceptor development and maintenance^36,41,42^. In *Crx-knockout* mice, photoreceptor cells failed to gain appropriate active chromatin configuration at *cis-regulatory regions* of CRX-dependent genes, including *PPR, RER and CBR* of the *Rho* locus^43^, implicating the role of these regions in epigenomic remodeling of *Rho* expression.

*In vivo* chromosome conformation capture assays (3C) have illustrated CRX/NRL-dependent intrachromasomal loop interactions between *RER* and *Rho PPR*/gene body^44^. These findings collectively suggest that *CBR* and *RER* are *Rho* enhancers, which potentially act by mediating local epigenomic changes to increase *PPR* activity. However, it is unknown if such enhancer-promoter interactions are necessary for *Rho* expression *in vivo*. More specifically, it remains to be determined if these two previously identified *Rho* enhancers are essential for the epigenomic modulation and transcriptional activation of the *Rho* gene, and what degree of regulation each contributes.

In order to fill this knowledge gap, we aimed to dissect and determine *in vivo* functional roles of the *Rho* enhancers (*CBR* and *RER*) and promoter (*PPR*) using a loss-of-function approach in mice. By delivering CRISPR/Cas9 and specific guide RNAs (gRNAs) to *C57BL/6J* mouse embryos, we generated mouse lines carrying <500bp deletions at each genomic region individually and a line carrying a paired deletion of both *CBR* and *RER*. We performed gene expression analyses and cellular phenotype characterization for each mouse line at various ages. Our results indicate that, as expected, the *Rho* promoter *PPR* is absolutely required for *Rho* expression, rod photoreceptor integrity and survival. However, to our surprise, the two enhancers (*CBR* and *RER*) collectively play only minor roles in *Rho* expression, which is only applicable in developing and aging retinas. *CBR/RER* deficiency did not largely impact rod development and maintenance up to one year of age. Together, our results suggest that *Rho* proximal promoter is necessary and sufficient for *Rho* expression in rod photoreceptors, while *Rho* enhancers are largely dispensable. Our findings highlight the importance of investigating the function of *cis-regulatory regions* in the native genomic context.

## RESULTS

### Generation of *Rhodopsin* (*Rho*) promoter and enhancer knockout mice

In order to examine the necessity of the *Rho* promoter and enhancers in *Rho* transcription, 4 mouse lines carrying a knockout of each genomic region were generated using CRISPR-Cas9/gRNA mediated excision. Based on the locations of CRX-ChIP-seq peaks ^39^, a pair of gRNAs targeting each peak at *Rho* promoter and two enhancer regions were introduced together with Cas9 enzyme into *C57BL/6J* mouse embryos to generate a 200-400bp deletion at each region. A mouse line lacking both enhancers (*CBR-/-RER-/-*) on the same allele was made using a sequential targeting approach, i.e. *RER* deletion on the *CBR-/-* allele by introducing *RER* gRNAs and Cas9 enzymes to the *CBR-/-* embryos. PCR combined with sanger sequencing confirmed the absence of each region with the indicated length of deletion: 217 for *PPR-/-* (retaining the TATA box and transcription start site), 234bp for *RER-/-* and 491bp for *CBR-/-* (Figure 1A). Mouse lines homozygous for each deletion appeared healthy and reproductive.

**Figure 1.**
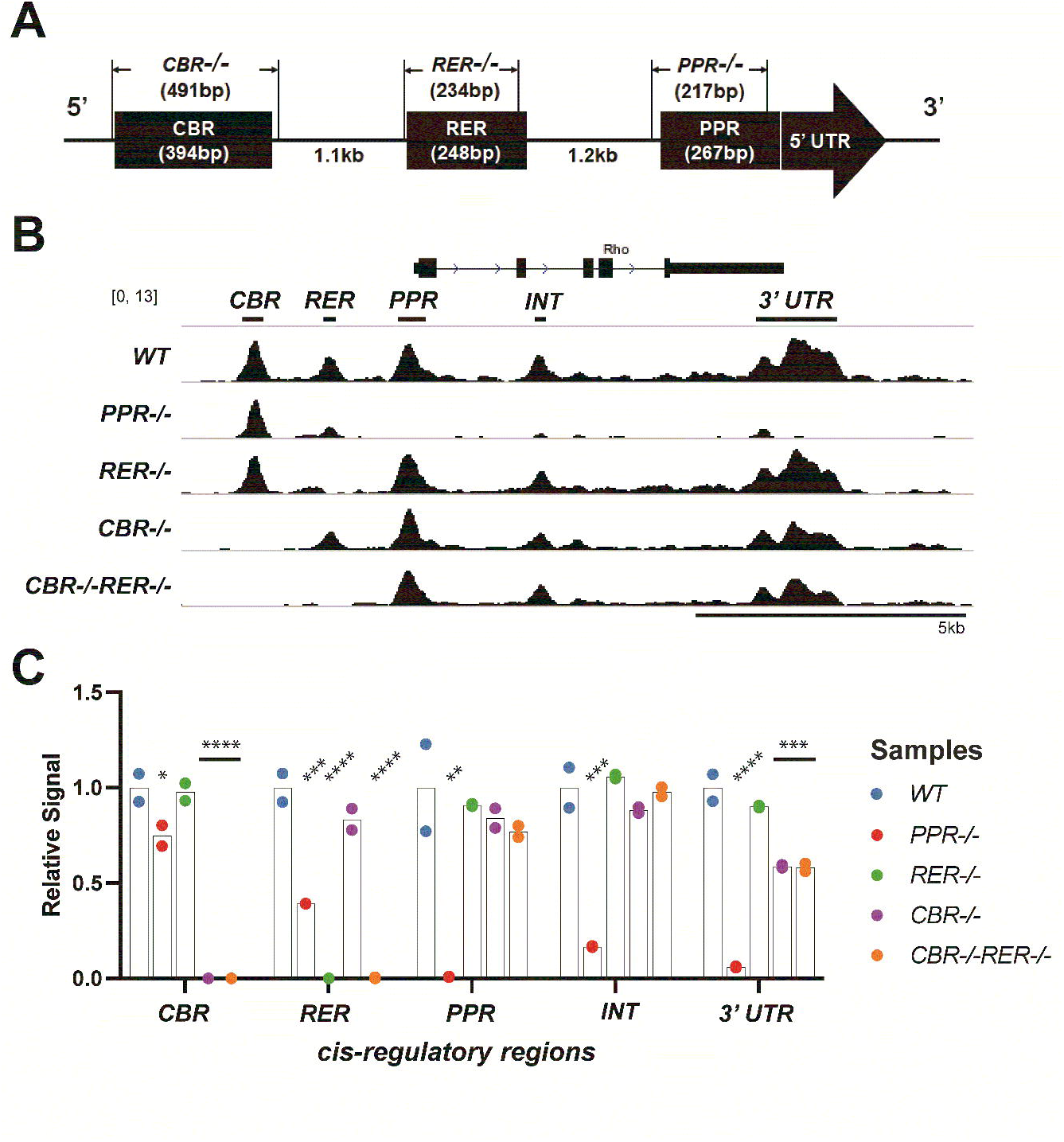
Changes of chromatin accessibility at *Rho* locus in mutant retinas. (**A**) Drawing of the *Rho* cis-regulatory regions (as black boxes) identified by previous studies and the specific knockouts involved in this study. (**B**) Browser tracks (mm10) of average ATAC-seq reads from duplicate retinal samples of the indicated genotypes at P14. Names of *Rho CREs* and peak calls are shown at the top of tracks. Scale bar represents 5kb. (**C**) Bar plot of average ATAC-seq signals relative to the *WT* control. One-way ANOVA test is performed. n=2 for each genotype. *CBR, CRX-bound region 1. RER, Rhodopsin enhancer region. PPR, Rhodopsin proximal promoter region. INT, Rhodopsin Intragenic region. 3’ UTR, 3’ untranslated region*. Asterisks (*, **, ***, ****) denote *p* ≤ 0.05, *p* ≤ 0.01, *p* ≤ 0.001, *p* ≤ 0.0001, respectively.

### Deletion of *Rho* promoter, but not enhancers, alters local chromatin accessibility

Since the chromatin of *Rho cis*-regulatory regions and gene body are activated during postnatal rod photoreceptor differentiation^45-47^, we tested if removal of *Rho* promoter or enhancers altered the chromatin accessibility of the *Rho* locus at postnatal day 14 (P14) using Assay for Transposase-Accessible Chromatin sequencing (ATAC-seq). In wildtype (*WT*) retina, rod photoreceptor specification is completed by P14 and the *Rho* gene is fully expressed; this timing is consistent with epigenetic remodeling of the *Rho* locus measured by ATAC-seq and DNAse I hypersensitivity profiling^47-49^. Duplicate ATAC-seq experiments on P14 whole retinas of *WT* and each of the four knockout mouse lines, *PPR-/-, CBR*-/-, *RER*-/-, and *CBR*-/-*RER*-/- yielded highly reproducible results. Consistent with the published results, *WT* samples displayed five significant open chromatin regions within the roughly 12kb region spanning the *Rho* locus (Figure 1B, *WT*), including the peaks at *CBR, RER, PPR*, and two intragenic regions (*INT* and *3’UTR*). The *PPR*-/- retina lost ATAC-seq signal not only at *Rho* promoter as expected, but also at *INT* and *3’UTR* (Figure 1B & 1C *PPR*-/-). Furthermore, *PPR* removal also caused a significant reduction of ATAC-seq signal upstream of the promoter, though the reduction was more profound at the proximal *RER* than at the more distal *CBR* (Figure 1B & 1C). These results suggest that *PPR* plays an essential role in achieving appropriate open chromatin configuration within *Rho* gene body, and is necessary for normal DNA accessibility at upstream enhancers *RER* and *CBR*. This role of *PPR* is specific to the *Rho* locus, as no significant DNA accessibility changes were detected at other loci in *PPR-/-* retina (Supplemental Figure 1A, Supplemental Figure 2A). Next, we analyzed changes in ATAC-seq signal in retinas of enhancer knockout lines. Interestingly, the removal of *RER* yielded no significant effect on any other regions, and the removal of *CBR* only affected the accessibility at the *3’UTR* (Figure 1B & 1C). Thus, neither *CBR* nor *RER* is required to establish the normal open chromatin configuration of the *Rho* locus during photoreceptor differentiation. Surprisingly, the removal of both *RER* and *CBR* did not produce an additive effect, but showed a similar impact as *CBR* single knockout (Figure 1B & 1C, *CBR-/-RER-/-*). Similar to the case of *PPR-/-*, global changes in chromatin accessibility were not observed in single- or double-enhancer knockout samples (Supplemental Figure 1B-D, Supplemental Figure 2A). Overall, these results suggest that *Rho* enhancers have limited functions in establishing *Rho* chromatin accessibility, but *PPR* plays an essential and enhancer-independent role.

Histone modifications can also be used as a readout of the local chromatin and transcriptional activation state. Both H3K4me3 and H3K27ac are associated with transcriptionally active chromatin. H3K4me3 usually marks active gene promoters, while H3K27ac is enriched at active enhancers and promoters ^5,9,50,51^. Both marks are detected at the actively transcribed *Rho* locus in mature mouse retina, which is defined as an active epigenetic state established in a CRX-dependent manner during postnatal development ^47,49^. In order to assess changes in histone marks upon *Rho* promoter or enhancer knockout, Cleavage Under Targets and Tagmentation (CUT&Tag) analysis was performed to report the high-resolution chromatin profiling on P14 retinal samples. Two validated antibodies specific to these histone marks were used in CUT&Tag experiments. In *WT* retina, H3K4me3 peaks showed a broad domain encompassing the gene body, starting at the *PPR* and extending to the final exon of *Rho* gene (Figure 2A, *WT*), but the signal was absent at *CBR* and *RER* regions. Analysis of mutants showed strikingly different landscapes: *PPR-/-* retina lost H3K4me3 deposition at the *Rho* locus while all enhancer-knockout samples showed comparable H3K4me3 marks to the *WT* control (Figure 2A, *PPR-/-*). Inspection of the 50kb region surrounding *Rho* confirmed that no other H3K4me3 sites were affected in the mutants (Supplemental Figure 2B). The results of H3K4me3 CUT&Tag analysis suggest that *Rho* enhancers neither contribute to H3K4me3 deposition nor affect *PPR* activity, while *PPR* is required for recruiting H3K4me3 to the *Rho* locus.

**Figure 2.**
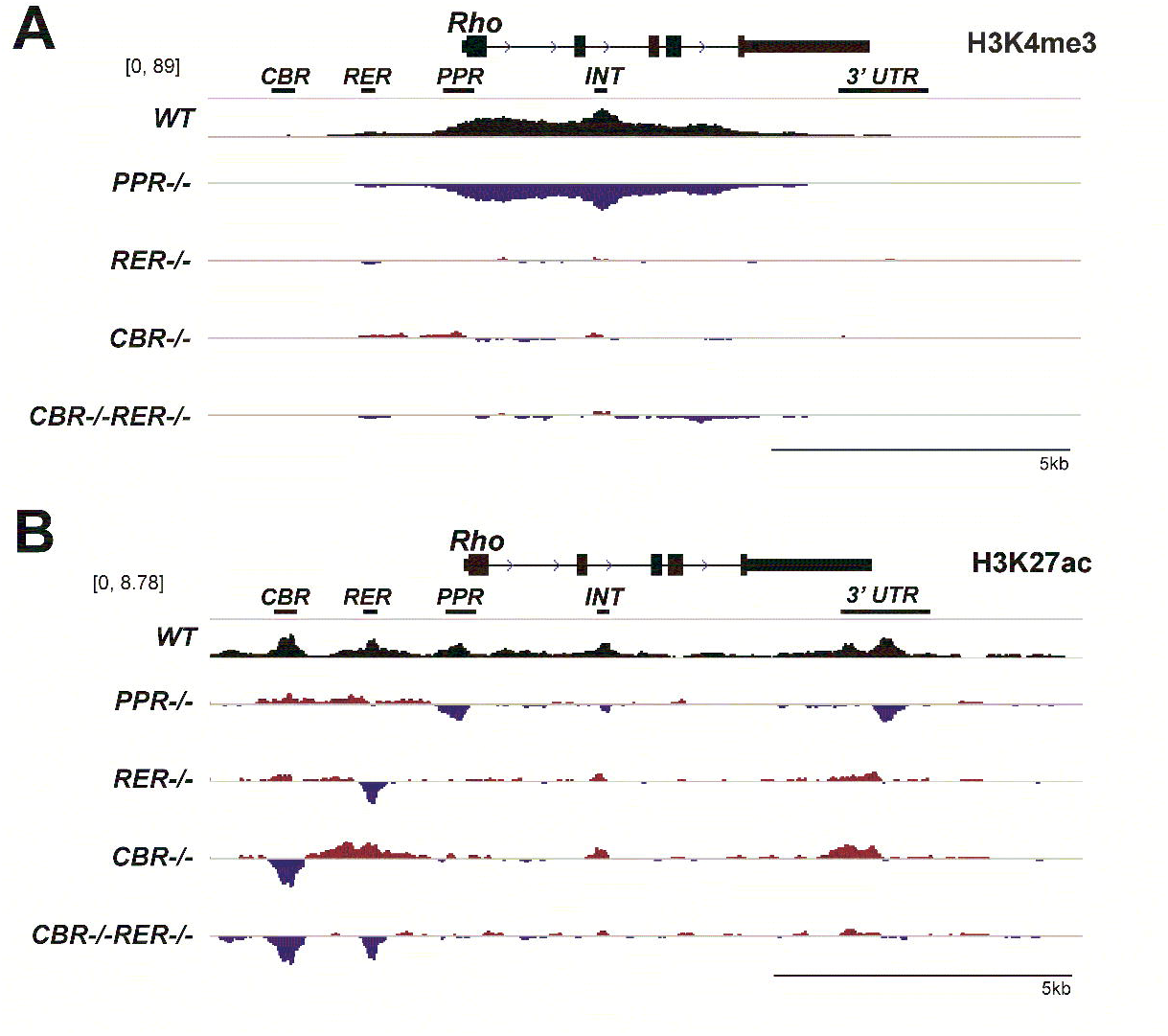
Changes of active histone marks, H3K4me3 and H3K27ac, at *Rho* locus in mutant retinas. Browser tracks display comparative analysis of H3K4me3 (**A**) and H3K27ac (**B**) peaks from retinal samples of the indicated genotypes at P14. Top track of each panel shows peaks in the *WT* control. The difference in H3K4me3 peaks ranges between -89 and 89. The difference in H3K27ac peaks ranges between -6.7 and 6.7. The following four tracks of each panel display increased (red) and decreased (blue) signals relative to the *WT* control in mutant retinas. Scale bar represents 5kb.

H3K27ac CUT&Tag analysis displayed 5 distinct peaks at *Rho* locus in *WT* retina at P14, including *CBR, RER, PPR, INT*, and *3’UTR*, respectively (Figure 2B, *WT*). In *PPR-/-* retina, H3K27ac signal was missing or greatly reduced at *PPR, INT* and *3’UTR*, but retained *WT* levels at *CBR* and *RER*, implying *CBR* and *RER* are capable of loading H3K27ac marks in the absence of *PPR* (Figure 2B, *PPR-/-*). In *RER*-/- retina, H3K27ac signal was similar to the *WT* control, besides a drop at the deleted *RER* region (Figure 2B, *RER-/-*). However, in *CBR-/-* retina, a compensatory increase in H3K27ac signal was observed at *RER, INT* and *3’UTR*, as well as at a few regions outside the *Rho* locus (Figure 2B, Supplemental Figure 2C, *CBR-/-*), possibly suggesting coordination or compensation between enhancers. In *CBR-/-RER-/-* retina, no change in H3K27ac signal was found at *PPR*, suggesting that the epigenetic activation state of the *Rho* promoter is unaltered by *CBR* and *RER* knockouts (Figure 2B, *CBR-/-RER-/-*). Collectively, H3K27ac CUT&Tag results suggest that *Rho* promoter and enhancers can independently modify their histone tails to establish an active epigenetic environment and there is little cross-talk between these three regulatory regions in establishing the local epigenetic state. Altogether, these CUT&Tag data indicate that *PPR* is absolutely required for establishing the chromatin configuration and active histone marks at the accessible regions of the *Rho* locus, especially at the promoter and intragenic regions, but *Rho* enhancers are largely dispensable for this remodeling.

### *Rho* enhancers make small and age-dependent contributions to *Rho* transcription levels

In order to analyze the effects of promoter/enhancer deletions on *Rho* expression, we performed next-generation RNA sequencing (RNA-seq) on P14 retinal samples of mutant and *WT* mice. Principal component analysis (PCA) showed that *PPR* samples were transcriptionally distinct from all other samples, evidenced by their distinct clustering (Figure 3A). *CBR*-/-*RER*-/- and *CBR-/-* samples were also separated from the *WT* controls (Figure 3A). Gene expression analysis showed that only *Rho* was differentially expressed in *PPR*-/- retina, and all other photoreceptor-enriched genes such as *Crx, Nrl, Nr2e3, Gnat1, Pde6a*, were not affected (Figure 3B). Analysis of other mutants showed similar profiles to the *WT* control, and *Rho* showed some degree of differential expression in *CBR*-/-*RER*-/- retina (Figure 3C). These results suggest that *PPR* plays a more dominant role in *Rho* transcription than either enhancer, and the combinatorial knockout of both enhancers causes only a minor decrease of *Rho* transcription.

**Figure 3.**
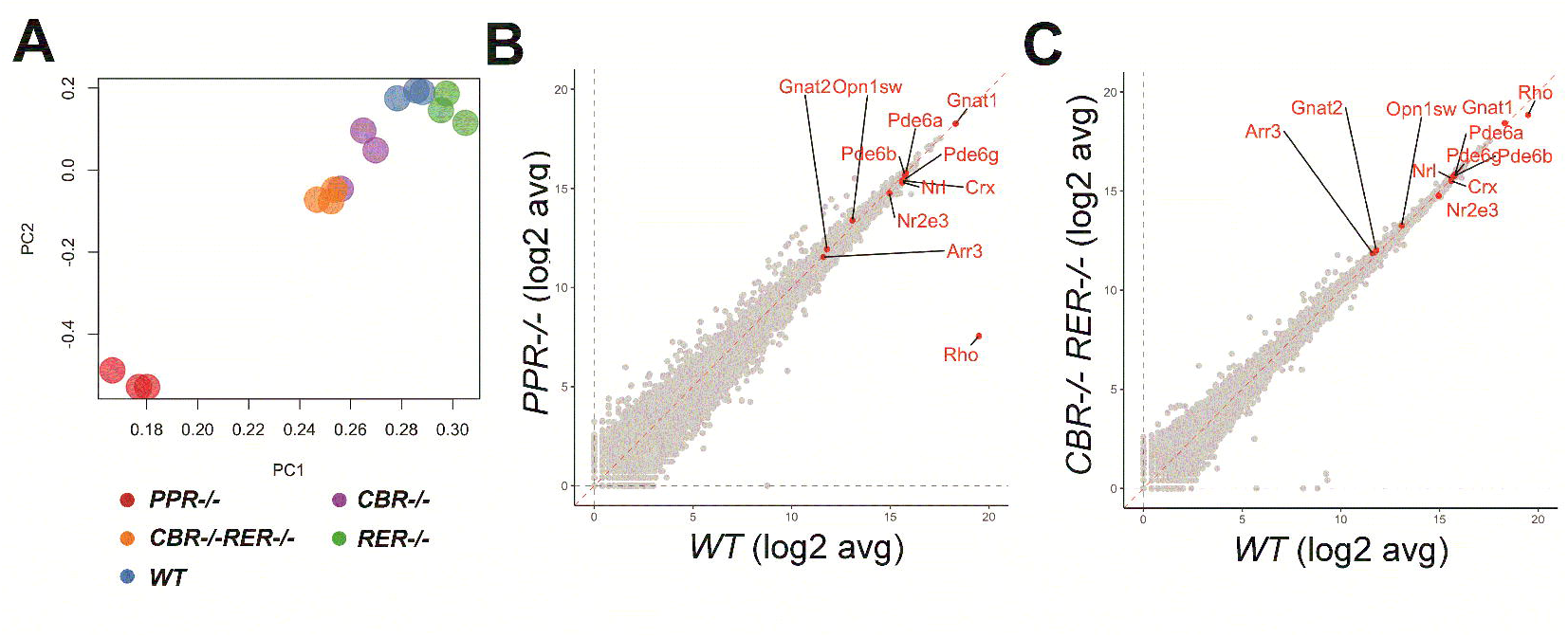
RNA-seq analysis of P14 mutant retinas. (**A**) PCA plot shows clusters of *WT* and the indicated mutant samples. (**B & C**) Scatterplot displays expression distributions of selected genes in the indicated mutant relative to the *WT* level. n=3 for each genotype.

We further investigated the degree by which promoter and enhancer knockouts impacted *Rho* expression using qRT-PCR with retinal samples of specific allelic arrangements at various ages. P14 retinal samples were collected for qRT-PCR experiments from genotypes illustrated in Figure 4A. At P14, *Rho* expression was completely ablated in *PPR*-/- retina and roughly reduced by half in *PPR+/-* retina (Figure 4B, *PPR-/-, PPR+/-*), suggesting that no compensatory regulation or feedback from the intact allele when one *PPR* allele is lost. In contrast, *RER-/-* retina had comparable *Rho* expression to the *WT* control, while *CBR-/-* retina reduced *Rho* expression to roughly 70-80% of the *WT* level (Figure 4B, *RER-/-, CBR-/-*), suggesting that *CBR*, not *RER*, makes a small contribution to *Rho* expression at P14. *CBR-/-RER-/-* retina showed reduced *Rho* expression to approximately 70-80% of the *WT* level, similar to, or slightly lower than that of *CBR*-/- retina (Figure 4B, *CBR-/-RER-/-*). These results suggest that the regulatory capacity of *Rho* enhancers, specifically *CBR*, contributes 20-30% of *Rho* expression. When a copy of *RER* was present in *Rho* enhancer regions (*CBR-/-RER+/-*), *Rho* expression generally matched with those in *CBR-/-* and *CBR-/-RER-/-* retinas, collectively suggesting that *RER* is not required for *Rho* expression (Figure 4B, *CBR-/-RER+/-*). When a copy of *CBR* was present in *Rho* enhancer regions (*CBR+/-RER-/-*), *Rho* expression reached to an indistinguishable level as the *WT* control (Figure 4B, *CBR+/-RER-/-*), suggesting that at least one copy of *CBR* is needed to achieve maximum-level *Rho* expression in the presence of two *PPR* copies. Lastly, *Rho* expression in a retina heterozygous for *PPR* and *Rho* enhancers was comparable to that of *PPR+/-* retina, i.e. about 50% to the *WT* control (Figure 4B, *CBR+/-RER+/-, PPR*+/-). These results together indicate that *PPR* is absolutely required for *Rho* expression and the regulatory functions of *Rho* enhancers are detectable and secondary to *PPR* in retinal development.

**Figure 4.**
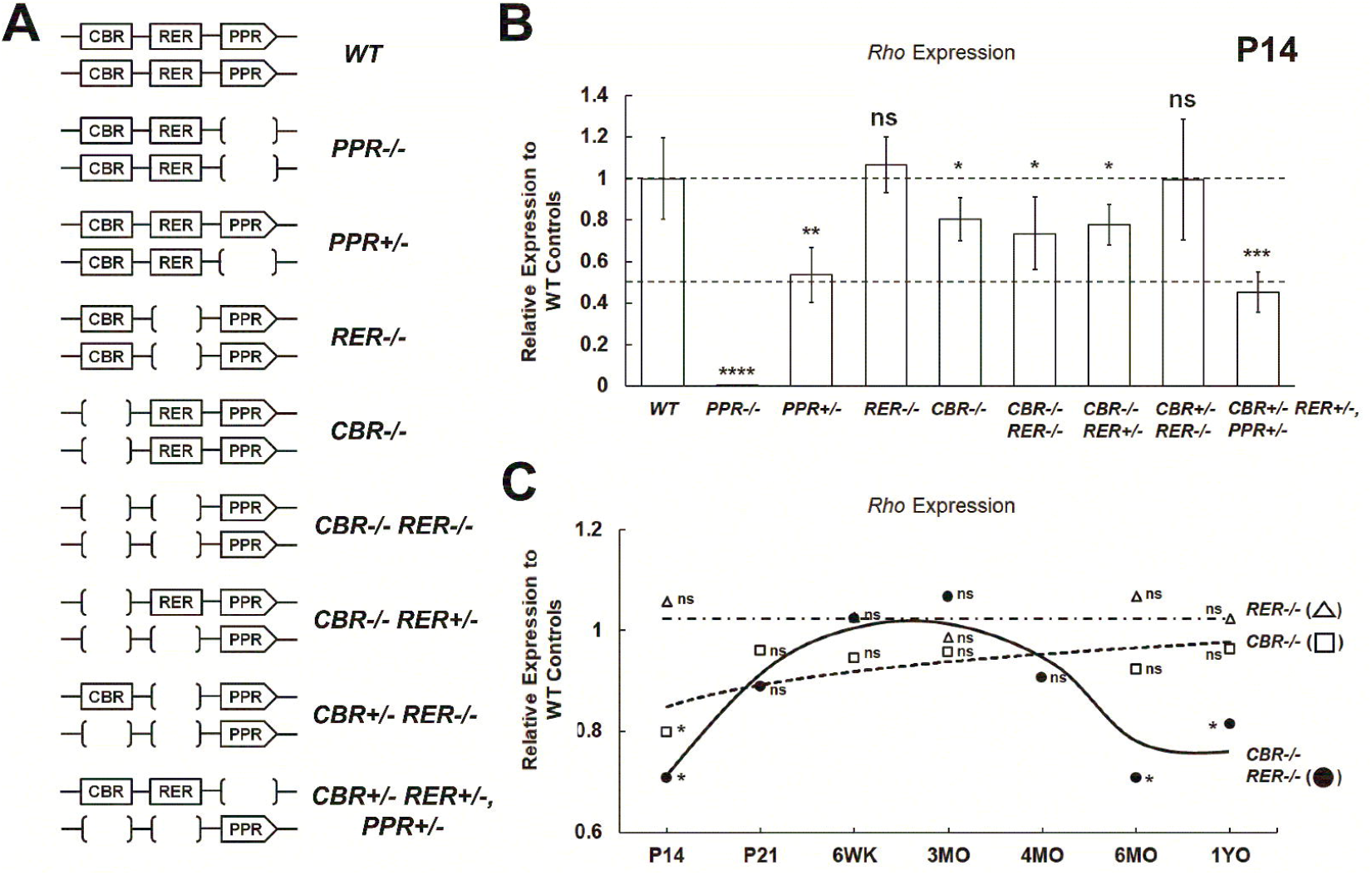
Quantitative PCR analysis of *Rho* expression in mutant retinas at various ages. (**A**) Diagrams of *WT* and mutant samples measured in this study. (**B**) qPCR analysis of *Rho* mRNA expression in P14 *WT* (n=8) and mutant retinas (n=10 for *CBR-/-*; *RER-/-*; *CBR-/-RER-/-*; n=8 for *CBR-/-RER+/-*; *CBR+/-RER-/-*; n=6 for *PPR-/-*; *PRR+/-*; *CBR+/-RER+/-, PPR+/-*). (**C**) Relative *Rho* expression profiles of *CBR-/-* (□), *RER-/-* (Δ), and *CBR-/-RER-/-* (●) samples up to 1YO (n ≥ 6 for each age). Interpolation is performed with MATLAB and Wolfram Mathematica: dashed line represents the *CBR-/-* profile, dash-dotted line represents the *RER-/-* profile, solid black line represents the *CBR-/-RER-/-* profile. All results are plotted as relative expression to *WT* control. Asterisks (*, **, ***, ****) denote *p* ≤ 0.05, *p* ≤ 0.01, *p* ≤ 0.001, *p* ≤ 0.0001, respectively by one-way ANOVA with Tukey’s multiple comparisons. ns means not significant.

Furthermore, we tested if *Rho* enhancers functioned to maintain *Rho* expression in adult retina. qRT-PCR experiments were performed with enhancer-knockout retinas up to 1 year-old (1YO). *Rho* expression in *RER*-/- retina always remained indistinguishable from that in the *WT* control (Figure 4C, *RER-/-*). *Rho* expression in *CBR-/-* retina initially reached about 70-80% of the *WT* level at P14 but increased steadily to a comparable level after the completion of retinal development and throughout adulthood (Figure 4C, *CBR*-/-). Interestingly, although *Rho* expression in *CBR-/-RER-/-* retina was as low as 70-80% to the *WT* control at P14, the relative *Rho* expression profile became comparable from P21 to 4 month-old (MO). However, at 6MO, the relative *Rho* expression dropped back to 70-80% of *WT* expression and persisted to 1YO (Figure 4C, *CBR*-/-*RER*-/-). Overall, these results suggest that *Rho* enhancers possess age-dependent regulatory activity to control *Rho* expression during rod differentiation and aging, but this regulation is only responsible for 20-30% of the overall expression. Besides changes in *Rho* expression in the *PPR*-/-, *PPR*+/- and *CBR*-/-*RER*-/- mutants, other rod photoreceptor-related genes such as *Crx* and *Gnat1* displayed comparable expression to the *WT* control at P14 (Supplemental Figure 4A) and 6MO (Supplemental Figure 4C). The immunohistochemistry (IHC) staining with anti-Rho and Gnat1 antibodies showed that both proteins were made in *CBR*-/-*RER*-/- and *PPR*+/- retinas and appeared at normal subcellular locations (Supplemental Figure 4B & 4D).

### Redundant roles of *Rho* enhancers contribute to the maintenance of rod morphology and function in old adults

We next investigated if the decrease in *Rho* expression induced any phenotypic changes in *Rho* enhancer and promoter mutants. *PPR*-/- retina failed to elaborate OS during development and showed only one remaining layer of cells within ONL at 6MO by H&E staining of retinal cross-sections (Figure 5A, *PPR*-/-), while *CBR*-/-*RER*-/- and *PPR*+/- retinas maintained well-laminated structures (Figure 5A, *CBR*-/-*RER*-/-, *PPR*+/-). No mislocalized cells were observed in the mutants. OS length of *CBR*-/-*RER*-/- and *PPR*+/- retinas appeared shorter than that of the *WT* control (Figure 4B, OS Length). ONL thickness in the *PPR+/-* retina was reduced as compared to *WT* control (Figure 5B, ONL Thickness), while *CBR*-/-*RER*-/- retina had a comparable ONL thickness. Since rod photoreceptors represent the most abundant cell type within ONL, these morphological measurements suggest that *CBR*-/-*RER*-/- retina very likely has indistinguishable cell numbers of rod photoreceptors to the *WT* control without apparent cell death (data not shown). Therefore, the slightly shorter OS of *CBR*-/-*RER*-/- retina is associated with reduced *Rho* expression. On the other hand, the decreased ONL thickness of *PPR+/-* retina indicates loss of rod photoreceptors. Furthermore, visual function of mutant retinas was assayed by dark-adapted (rod response) electroretinography (ERG). As expected, *PPR-/-* mice had no ERG responses, while *PPR+/-* mice had decreased dark-adapted A-wave and B-wave amplitudes at high-light intensities as compared to the *WT* control (Figure 5C). *CBR*-/-*RER*-/- mice also showed lower A-wave and B-wave amplitudes at the high light intensities, but less severe than *PPR+/-* (Figure 5C). This is consistent with the reduction of *Rho* expression and OS length in the *CBR*-/-*RER*-/- retina that was less severe than observed in the *PPR*+/- retina. In addition, 6MO *CBR*-/- and *RER*-/- single mutants were indistinguishable from the *WT* control in retinal morphology (Supplemental Figure 5A), OS length (Supplemental Figure 5B) and ERG responses (Supplemental Figure 5C & 5D), echoing their comparable *Rho* expression as the *WT* control. Thus, *CBR* and *RER* act redundantly to constitute the *Rho* enhancer landscape for transcriptional regulation in adult retina, and their combined loss leads to measurable functional deficits.

**Figure 5.**
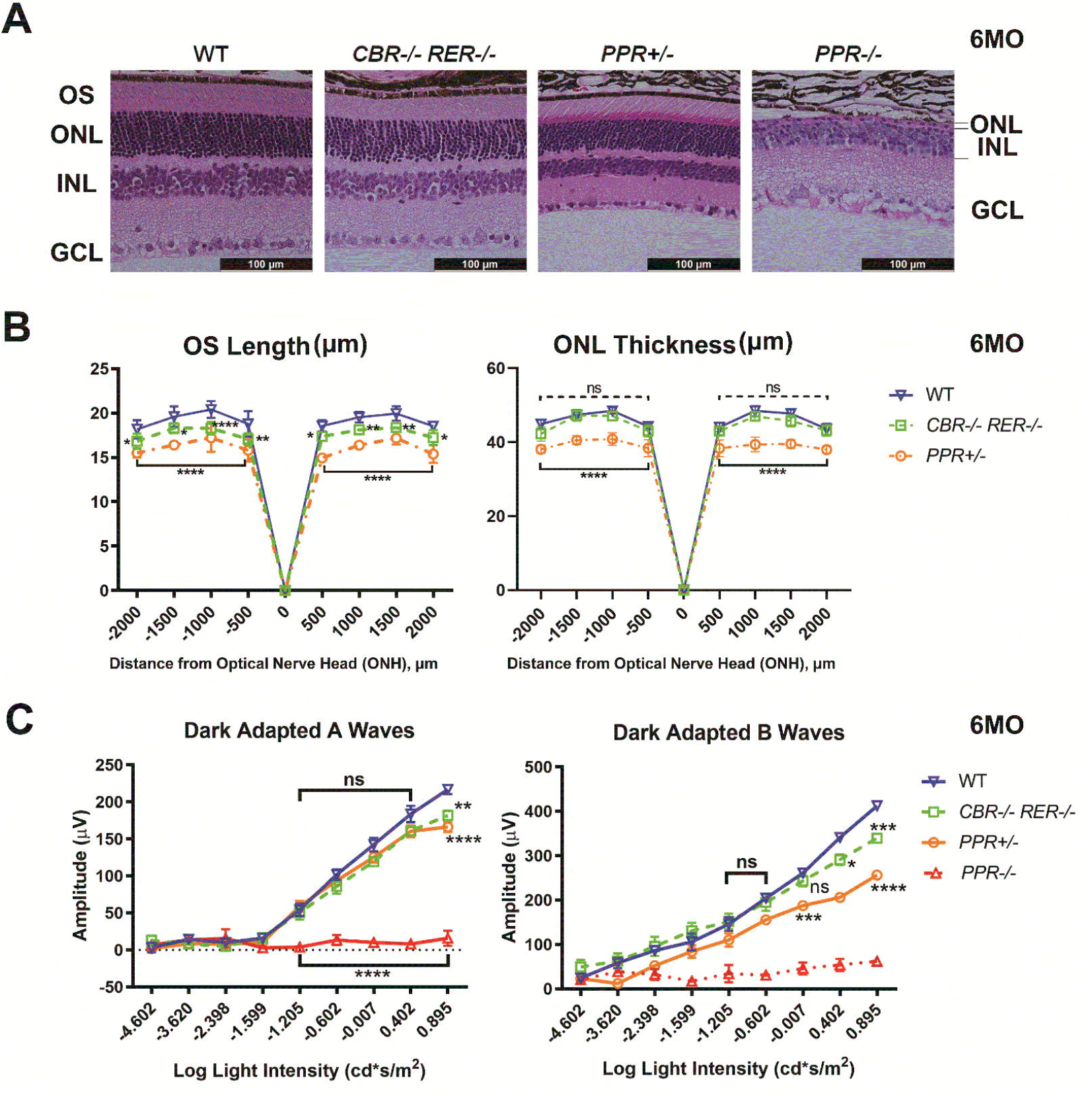
Changes of retinal morphology and function in mutants at 6MO. (**A**) Hematoxylin and Eosin (H&E) cross-section staining of 6MO *WT* and mutant retinas of the indicated genotypes. OS: outer segment; ONL: outer nuclear layer; INL: inner nuclear layer; GCL: ganglion cell layer. Scale bar represents 100 μm for all image panels. (**B**) OS thickness (left panel, in μm) and ONL thickness (right panel, in μm) in 6MO *WT* and mutant retinas of the indicated genotypes at various positions from the optic nerve head (ONH). Error bars represent mean (SD) (n ≥ 5). Dashed black line represents the comparison between *WT* and *CBR-/-RER-/-* samples. Solid black line represents the comparison between *WT* and *PPR+/-* samples. (**C**) Electroretinogram (ERG) analysis of 6MO *WT* and mutant mice of the indicated genotypes, showing amplitude changes of dark-adapted A-waves (left panel) and B-waves (right panel). Mean amplitudes (μV) are plotted against stimulus light intensity. Error bars represent SEM (n ≥ 7). All statistics is done by comparing to *WT* control. Asterisks (*, **, ***, ****) denote *p* ≤ 0.05, *p* ≤ 0.01, *p* ≤ 0.001, *p* ≤ 0.0001, respectively by two-way ANOVA with Tukey’s multiple comparisons. ns means not significant.

Rod morphological and functional changes in mutant retinas were also assessed at early adult ages. At 6 weeks (6WK) and 3MO, OS length of the *PPR*+/- retina appeared shorter (Supplemental Figure 6A & 6B). This morphological defect in the *PPR*+/- retina echoed the lower amplitudes of dark-adapted ERG A-waves at the highest light intensity as compared to that of the *WT* control (Supplemental Figure 6E & 6F). OS length and the amplitudes of dark-adapted ERG A-wave of the *CBR*-/-*RER*-/- retina were comparable to *WT* controls (Supplemental Figure 6A, 6B, 6E & 6F). Importantly, ONL thickness of all mutants remained comparable to that of the *WT* control at 6WK and 3MO (Supplemental Figure 6C & 6D). Compared to the ONL reduction in 6MO *PPR+/-* retina, these results suggest that *PPR+/-* retinal degeneration occurs after 3MO. As a reference, OS and ONL measurements in heterozygous and homozygous *Rho*-*knockout* mice^31^ (*Rho-/-* and *Rho+/-*) showed similar results as *PPR* mutants: ONL thickness of *PPR*-/- retina degenerated dramatically after 6WK (Supplemental Figure 7A, *PPR-/-*). *PPR*+/- retina began to show ONL degeneration at 6MO but reduced OS length at 3MO, similar to that of *Rho+/-* retina (Supplemental Figure 7A & 7B, *PPR+/-* vs. *Rho+/-*).

## DISCUSSION

Using CRISPR-Cas9/gRNA mediated excision, this study examined the specificity and strength of two *Rho* enhancers, namely *CBR* and *RER*, and of *Rho* promoter in regulating *Rho* expression. Previous reporter-based assays showed that both the *RER* and *CBR* were capable of enhancing *Rho* promoter activity by 8-10 fold in postnatal mouse retina^33,39^. However, this study shows that knockout of each or both enhancers surprisingly produce null or only moderate impact (20-30% loss) on endogenous *Rho* expression. Notably, *RER*, that was previously identified by a study with transgenic mice^33^ and known to interact with *PPR* by 3C assay^44^, fails to display any necessity for endogenous *Rho* expression in developmental and adult ages. Consistent with their minor roles, single *Rho* enhancer knockout also shows limited impact on rod photoreceptor identify, health and function. These results are in contrast to the essential function of the *Locus Control Region* (*LCR*) in its regulation of red/green (R/G) cone opsin expression^52-54^. The conserved *LCR* is absolutely required for mutually exclusive expression of red vs green opsin in the respective cone subtype in humans, mouse and zebrafish models. *RER* and *R/G LCR* share a conserved core region of similar TF binding motifs, including CRX binding sites^33^. Several models might explain the minimal relevance of *Rho* enhancers *in situ*, but high activity in reporter assays: firstly, the strength of a promoter might determine its cooperation with enhancers. *Rho* promoter (*PPR*) is highly robust and sufficient to drive cell type-specific expression of target genes both in episomal vectors^39^ and at ectopic genomic locations^33^. Future studies are needed to determine the relative strength of *Rho* promoter vs other photoreceptor gene promoters and how the promoter strength influences functional interactions with specific enhancers. Secondly, dynamic expression and precision might also be a determinant. Genes coding for cell fate-determining transcription factors with complex spatial and temporal expression patterns often harbor strong distal enhancers or enhancer clusters, such as those found in *Pax6*^*55*^, *Otx2*^*56-58*^, *Vsx2*^*59*^, *Atoh7*^*60*^ *and Blimp1*^*61*^. *Rho* expression, despite being one of the highest expressed gene in the retina, follows a simple induction curve during rod photoreceptor development^62^, which may not require a prominent enhancer but a strong self-functioning promoter. Finally, results of reporter assays may have exaggerated *Rho* enhancer activities, as reporter gene constructs do not contain the correct genomic context, including native gene structure and epigenomic landscape. Overall, the findings of this study highlight the importance to validate functions of reporter-identified enhancers in a native genomic context.

This study shows age-dependent activities of *Rho* enhancers. During postnatal development, *CBR* clearly plays a dominant role, as it is responsible for 20-30% *Rho* expression, while *RER*’s activity was dispensable. Thus, *CBR*, but not *RER*, is required for appropriate *Rho* transcription during development. However, this minor contribution of *CBR* does not affect the development of rod subcellular structure and function. In adult retinas, deletion of *Rho* enhancers results in no detectable affects in young and mid ages, but after 6 MO, *RER/CBR* knockout causes a loss of 20-30% of *Rho* transcription. These suggest that *CBR* and *RER* play a redundant role in maintaining high-level *Rho* transcription in aged retinas. Similar “shadow” or redundant enhancers have been reported elsewhere, such as individual enhancers in *β-globin LCR, Pax6* lens enhancers and *Atoh7* distal enhancer ^55,60,63,64^. These enhancers mostly act during tissue/organ genesis and are required for regulating the expression timing of specific genes. Our discovery of redundant enhancers in aged retina provides a new insight into enhancers’ roles in aging and tissue-specific maintenance.

Consistently, *CBR/RER* knockout mutants affected *Rho* expression as well as morphological and functional integrity at 6MO. Since *PPR+/-* (50% *Rho* reduction) produced a more severe phenotype than *CBR/RER* knockout mutants (20-30% *Rho* reduction) in aged adults, our results confirm that rod homeostasis requires precise control of *Rho* expression. Aged rod photoreceptors with disrupted *Rho* enhancer activity may be more susceptible to genetic and/or environmental insults. Future studies are required to determine if *Rho* enhancer deficiency complicates (or modifies) defective phenotypes under disease/stress conditions, particularly in progressive rod photoreceptor degeneration.

Typical “super enhancer” characteristics were detected in wildtype retina within 5 kb of the *Rho* upstream regulatory sequence. Changes of these characteristics in mutants, in general, agreed with the impact of each region on *Rho* transcription. In particular, loss of ATAC signals at *Rho* gene body was only seen in *PPR-/-*, but not in enhancer knockout mutants, highlighting the interdependency and necessity of *PPR* in epigenomic modulation during *Rho* transcription activation. Deleting *CBR* and/or *RER* enhancers had limited impact on the epigenomic landscape of the *Rho* locus, despite the 20-30% contribution to transcription, suggesting that the two enhancers do not act in a “*trans”* manner on DNA accessibility or chromatin configuration. Most likely, they promote RNA polymerase II activity at the promoter and gene body via CRX/NRL-dependent looping interactions^44^. A similar interactive pattern of predominant promoter versus subsidiary enhancers has been previously reported^65^. Although *CBR* and *RER* deletions are insufficient to elicit large changes in *Rho* expression and DNA accessibility of *Rho PPR*, this study cannot exclude the existence or importance of more distal *Rho*-specific *CREs* in regulating chromatin dynamics at the *Rho PPR* region.

The mouse *Rho* gene shares about 95% sequence homology with the human gene^33^ (Mouse Genome Informatics). Importantly, *Rho* enhancers at the *Rho* locus also share remarkably conserved topography between the two species in terms of *CRE* number and relative locations^33,39^. The profile of transcription factors bound to *Rho* enhancers and promoter is also conserved^39,66^. While numerous non-synonymous and nonsense variants within the *Rho* coding region have been found to cause human retinal disease, no regulatory variants have yet been discovered. This study suggests that future efforts should be primarily focused on the proximal promoter region, particularly at critical TF binding motifs. Distal regulatory variants within the *RER* and *CBR* are unlikely to cause pathogenic transcriptional deficits for severe developmental defects. These variants however could be genetic modifiers of other pathogenic mechanisms or independently yield minor but detectable effects on visual function and alter long-term photoreceptor health.

In conclusion, this study has described unexpectedly minor roles of two previously identified *Rho* enhancers on *in vivo Rho* transcription, and their redundant activity in rod functional maintenance in aging retinas. These findings contribute to our understanding of enhancer functions and mechanism of action *in vivo*, and highlight the importance of dissecting individual enhancer functions and interactions in the native genomic context at various ages.

## MATERIALS AND METHODS

### Animals

All mice in this study were on the *C57BL/6J* background. Both male and female mice were used in experiments. All animal procedures were conducted according to the Guide for the Care and Use of Laboratory Animals of the National Institute of Health, and were approved by the Washington University in St. Louis Institutional Animal Care and Use Committee.

### Generation of mutant mice

CRISPR/Cas9 gene editing was performed using two single-guide RNA (sgRNAs) flanking each element of *CBR, RER, PPR* in order to delete these elements (see Figure 1). The gRNA sequences are: For CBR, 5’:TAGCTCCGTTTCCACATTGA and 3’:TCAAGATACACTGTCCCCAC. For RER, 5’: GCTTCATCGTGGTCTCCGCG and 3’: TCCATGCAGGTGTCTTGTTT. For *PPR*, 5’: GAAGTGAATTTAGGGCCCAA and 3’: GCGGATGCTGAATCAGCCTC. Fertilized mouse oocytes injected with sgRNAs were allowed to proceed to birth to generate mouse lines. In order to generate the mutant carrying deletions of both *CBR* and *RER* on the same chromosome, a mouse line homozygous for *CBR*-/- was used to prepare fertilized oocytes that were subsequently injected with sgRNAs for *RER* deletion. F0 mutants were genotyped by DNA sequencing of PCR products overlapping the mutations.

### Histology and immunohistochemistry

Eyes were enucleated at tested ages and fixed in 4% paraformaldehyde at 4°C overnight for paraffin embedded sections. Each retinal cross-section was cut 5 microns thick on a microtome. Hematoxylin and Eosin (H&E) staining was used to examine retinal morphology.

For IHC staining, sections firstly went through antigen retrieval with citrate buffer, and blocked with a blocking buffer of 5% donkey serum, 1% BSA, 0.1% Triton-x-100 in 1X PBS (pH-7.4) for 2 hours. Sections were then incubated with primary antibodies at 4°C overnight, then washed, followed by 2-hour incubation of specific secondary antibodies. Primary antibody to Rho (MilliporeSigma, O4886) or Gnat1 (MilliporeSigma, GT40562) and secondary antibodies were applied with optimal dilution ratios.

All slides were mounted with hard-set mounting medium with DAPI (Vectashield, Vector Laboratories, Inc., CA).

### Electroretinogram

ERGs were performed on adult mice at different ages using UTAS-E3000 Visual Electrodiagnostic System (LKC Technologies Inc., MD). Mice were dark-adapted overnight prior to the tests. The experimental procedures were previously established in our lab. ERG responses of biological replicates were recorded, averaged and analyzed using Graphpad Prism 8 (GraphPad Software, CA). The mean peak amplitudes of dark-adapted A and B waves and light-adapted B waves were plotted against log values of light intensities (cd*s/m^2^). The statistical analysis was done by two-way ANOVA with multiple pairwise comparisons (Tukey’s).

### Quantitative PCR

Each RNA sample was extracted from 2 retinae of a mouse using the NucleoSpin RNA Plus kit (Macherey-Nagel, PA). RNA concentrations were measured using a NanoDrop One spectrophotometer (ThermoFisher Scientific). 1 μg of RNA was used to synthesize cDNA with First Strand cDNA Synthesis kit (Roche, IN). The reaction master mix contained EvaGreen polymerase (Bio-Rad Laboratories, CA), 1 μM primer mix, and diluted cDNA samples. Samples were run using a two-step 40-cycle protocol on a Bio-Rad CFX96 Thermal Cycler (Bio-Rad Laboratories, CA). At least 3 technical triplicates were run for each gene. Primers (5’ to 3’) used in this study were *Rho* (F: GCTTCCCTACGCCAGTGTG, R: CAGTGGATTCTTGCCGCAG), *Crx* (F: GTCCCATACTCAAGTGCCC, R: TGCTGTTTCTGCTGCTGTCG), *Gnat1* (F: ACGATGGACCTAACACTTACGAGG, R: TGGAAAGGACGGTATTTGAGG). Data were analysed with QBase software (Biogazelle, Belgium). The statistical analysis was done by Student’s t-test with *p* < 0.05, CI:95%.

### RNA-seq data generation and analysis

RNA was extracted from retinal samples using the NucleoSpin® RNA kit (Macherey-Nagel) using the manufacturer’s protocol. Total RNA integrity was determined using Agilent Bioanalyzer or 4200 Tapestation. Library preparation and sequencing experiments were performed at Genome Technology Access Center, Washington University in St. Louis. A brief description of the experimental procedures was outline as follows. Library preparation was performed with 5 to 10ug of total RNA with a RIN score greater than 8.0. Ribosomal RNA was removed by poly-A selection using Oligo-dT beads (mRNA Direct kit, Life Technologies). mRNA was then fragmented in reverse transcriptase buffer and heating to 94 degrees for 8 minutes. mRNA was reverse transcribed to yield cDNA using SuperScript III RT enzyme (Life Technologies, per manufacturer’s instructions) and random hexamers. A second strand reaction was performed to yield ds-cDNA. cDNA was blunt ended, had an A base added to the 3’ ends, and then had Illumina sequencing adapters ligated to the ends. Ligated fragments were then amplified for 12-15 cycles using primers incorporating unique dual index tags. Fragments were sequenced on an Illumina NovaSeq-6000 using paired end reads extending 150 bases. Reads were trimmed using Trim-Galore! (v0.6.7) a wrapper for Cutadapt (v3.4) and FastQC (v0.11.9). Trimmed reads were mapped to mm10 using STAR (v2.7.0)^67^. Mapped reads were cleaned using Samtools (v1.9.4), gene counts generated by HTseq (v0.11.2)^68^, and visualizations generated in R (v3.6.1).

### ATAC-seq data generation

ATAC-seq libraries were generated as published in Ruzycki et al^49^, a slightly modified protocol to that published by Buenrostro et al^69^. Briefly, retinas were dissected from P14 mice of each genotype and washed in PBS. Retinal cells were dissociated using 2% collagenase in TESCA buffer for 13 minutes at 37C. DNase I (0.5 Units; Roche, Basel, Switzerland) was added for the final 3 min to minimize clumping. DMEM+10% FBS was added to stop the reaction. Nuclei were stained with SYBR Gold (Thermo Fisher Scientific) diluted 1:100 in trypan blue and counted using a fluorescent microscope. 50,000 cells were resuspended in TD buffer for a 1hr incubation at 37C with TDE1 (Nexetra DNA Library Prep Kit; Illumina, San Diego, CA). Remaining steps of library prep were performed as published by Buenrostro et al. Final libraries were pooled and sequenced using the Illumina 2500.

### CUT&Tag data generation

Retinal cells were dissociated and counted in same manner as described above for ATAC-seq. After quantification, cells were diluted, aliquoted and cryopreserved by adding DMSO to a final volumne of 10% and slowly freezing to –80C in a Mr. Frosty container. CUT&Tag reactions were performed using Epicypher reagents (protocol v1.5; Epicypher, USA). Briefly, cells were thawed on ice and 100,000 bound to ConA beads for each sample. Primary antibody incubation was left at 4C overnight on a nutator. 0.5ug of antibody was used for each reaction; H3K4me3 (MABE647; EMD Millipore Corporation, USA) and H3K27Ac (ab177178; Abcam Inc, USA). 0.5ug secondary antibody (CUTANA anti-rabbit secondary antibody; 13-0047; Epicypher) was incubated for 0.5hr at RT, and Tn5 binding and trnasposition done exactly as specified in CUTANA protocol. After transposition, custom 10bp unique-dual-indexed primers were used to amplify transposed fragments (P7-CAAGCAGAAGACGGCATACGAGAT-10bpBC-GTCTCGTGGGCTCGGAGATGTG, P5-AATGATACGGCGACCACCGAGATCTACAC-10bpBC-TCGTCGGCAGCGTCAGATGTGTAT) for 20 cycles. Libraries were cleaned and size-selected using Ampure XP beads. Libraries were pooled and sequenced using the Illumina NovaSeq 6000.

### ATAC-seq and CUT&Tag data analysis

Reads were trimmed using Trim-Galore! (v0.6.7) a wrapper for Cutadapt (v3.4) and FastQC (v0.11.9). Trimmed reads were mapped to the mm10 genome build using bowtie2 (v2.4.1)^70,71^. Mapped reads were cleaned using Samtools (v1.9.4), Picard (v2.25.7), and Bedtools (v2.27.1). For visualization, reads were converted to bigwig format and compared using Deeptools (v3.5.1). ATAC-seq peak calling was accomplished using MACS2 (v2.2.7.1).

### Data access

All raw and processed data has been uploaded to the NCBI SRA database.

## ETHICS STATEMENT

All animal procedures were conducted according to the Guide for the Care and Use of Laboratory Animals of the National Institute of Health, and were approved by the Washington University in St. Louis Institutional Animal Care and Use Committee.

## AUTHOR CONTRIBUTIONS

SC conceived of the study, CS, PR and SC designed the experiments, CS and PR performed experiments and analyzed the results. CS, PR and SC wrote the manuscript. All authors read and approved the final manuscript.

## ACKNOWLEDGMENTS

We thank Courtney Linne, Mingyan Yang, Guangyi Ling, Belinda Dana and Dr. Xiaodong Zhang for technical assistance; We also thank Susan Penrose and Mike Casey with the Molecular Genetics Service Core for generating *Rho* promoter/enhancer knockout mice. This work was supported by NIH grants EY012543 (to SC) and EY002687 (to WU-DOVS), and unrestricted funds from Research to Prevent Blindness (to WU-DOVS).

