## Supplemental Materials for "*Rho* enhancers play unexpectedly minor roles in *Rhodopsin* transcription and rod cell integrity"

Chi Sun[1], Philip A. Ruzycki[1,2]* and Shiming Chen[1,3]*
[1]Department of Ophthalmology and Visual Sciences, [2]Department of Genetics, and [3]Department of Developmental Biology, Washington University, Saint Louis, Missouri, USA

*To whom the correspondence should be addressed.
Shiming Chen
660 South Euclid Avenue, MSC 8096-0006-06
St. Louis, MO 63110, USA


Philip Ruzycki
660 South Euclid Avenue, MSC 8096-0006-11
St. Louis, MO 63110, USA


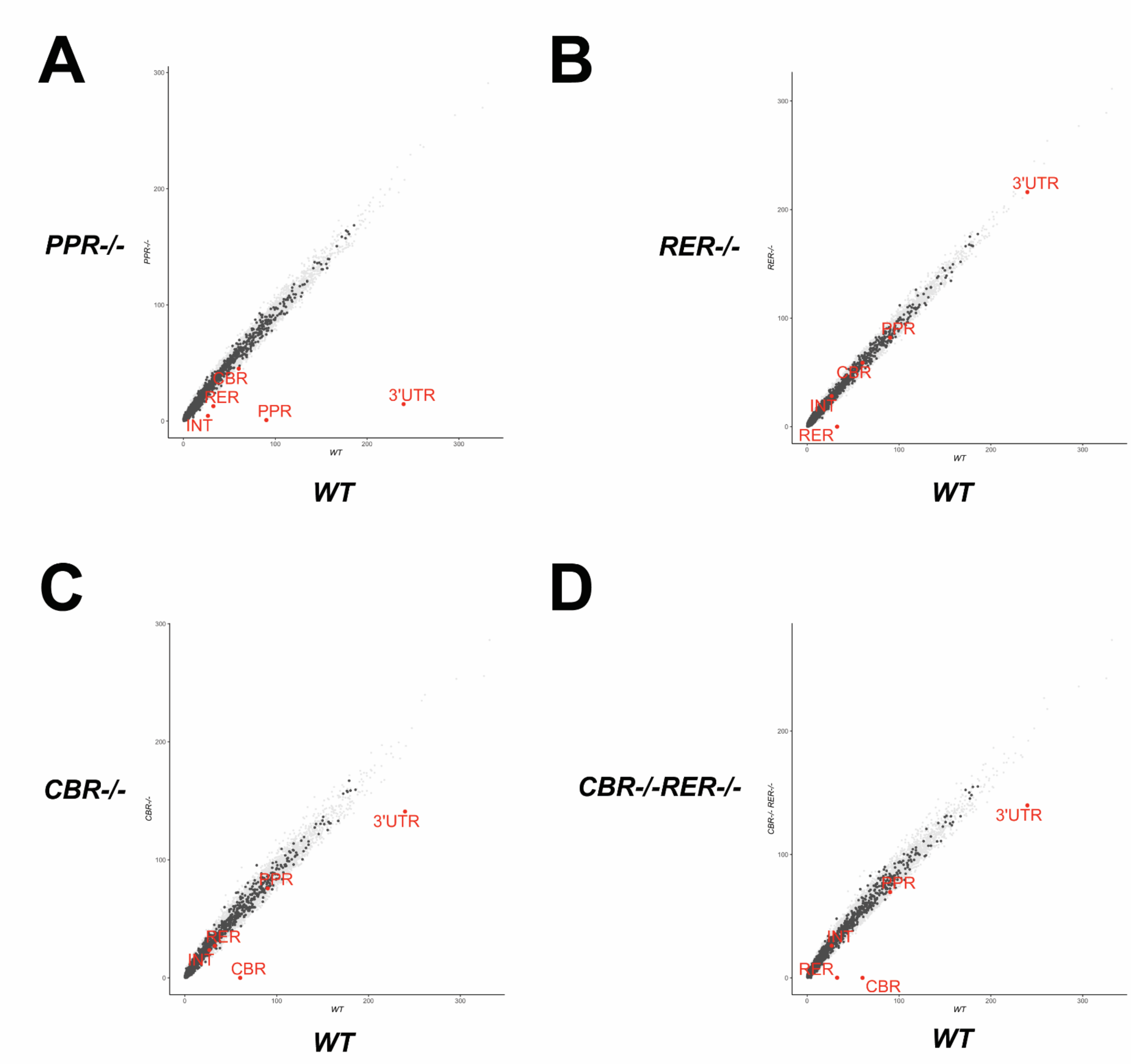


**Supplemental Figure 1**. **Scatterplot displays ATAC-seq signal changes at *Rho cis-regulatory regions* in the indicated mutants relative to the *WT* control**. *CBR*, *CRX-bound region 1*. *RER*, *Rhodopsin enhancer region*. *PPR*, *Rhodopsin proximal promoter region*. *INT*, *Rhodopsin Intragenic region*. *3' UTR*, *3' untranslated region*.

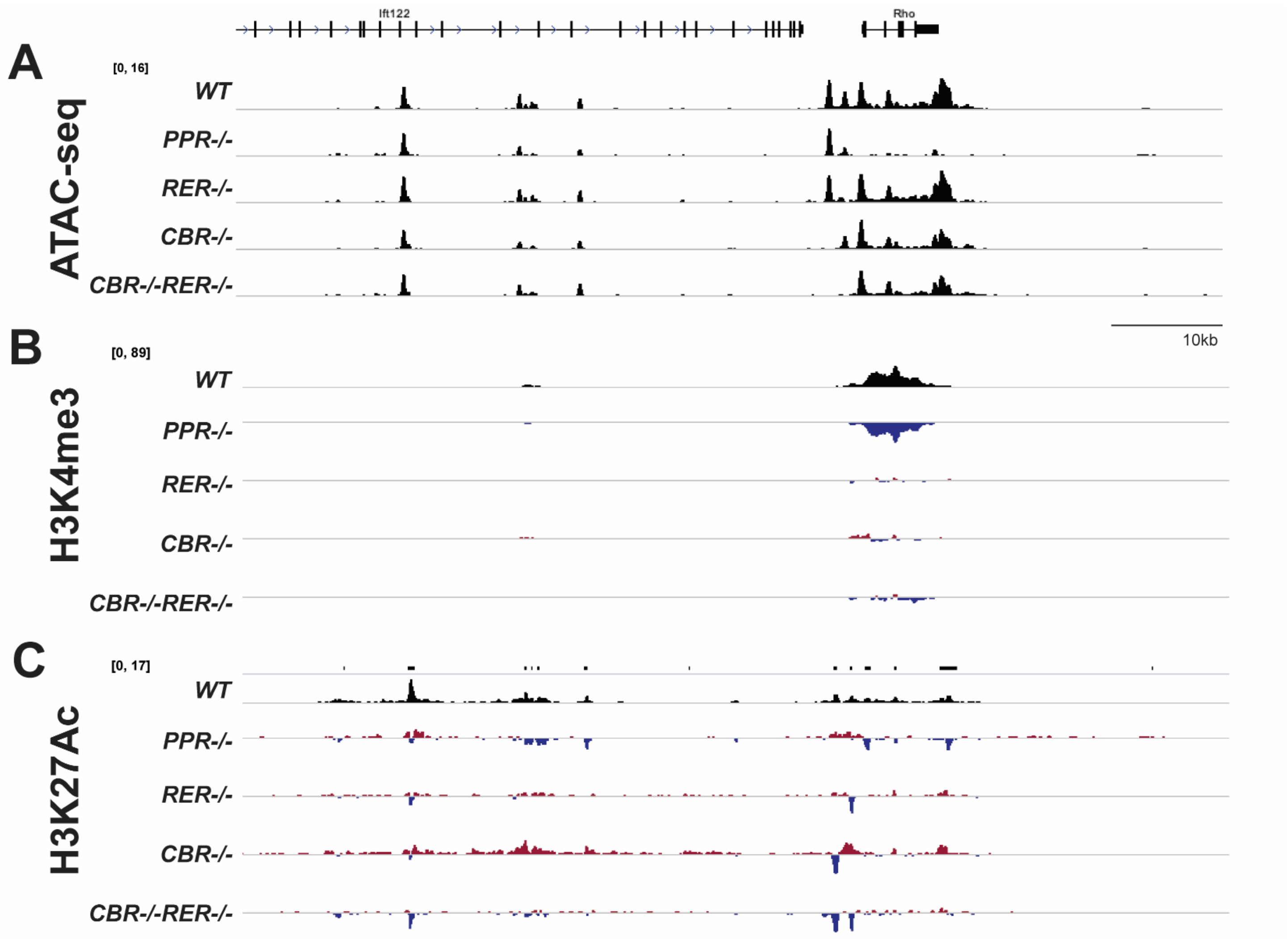


**Supplemental Figure 2. Browser tracks display ATAC-seq (A), H3K4me3 (B), H3K27ac (C) signals within 50 kb upstream of *Rho* locus.** A 10 kb bar is included for estimating size of the genomic region. The difference in H3K4me3 peaks ranges between -89 and 89. The difference in H3K27ac peaks ranges between -6.7 and 6.7.

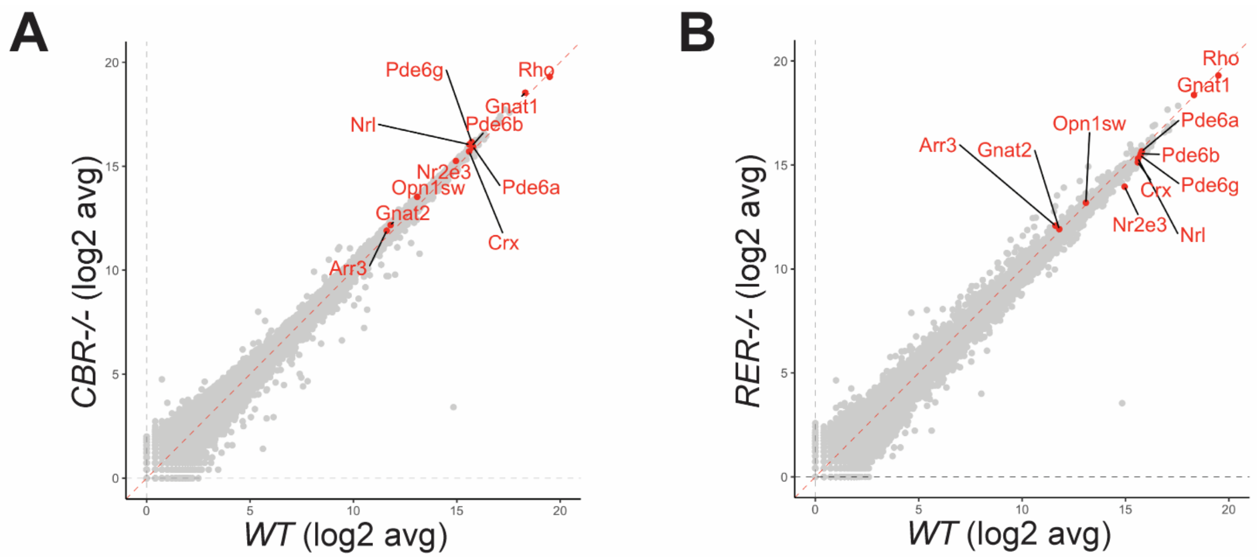


**Supplemental Figure 3. RNA-seq analysis in P14 *CBR-/-* (A) and *RER-/-* (B) retinas.** Each scatterplot displays the expression distribution of selected genes in the indicated mutant (Y-axis) relative to the *WT* control (X-axis).

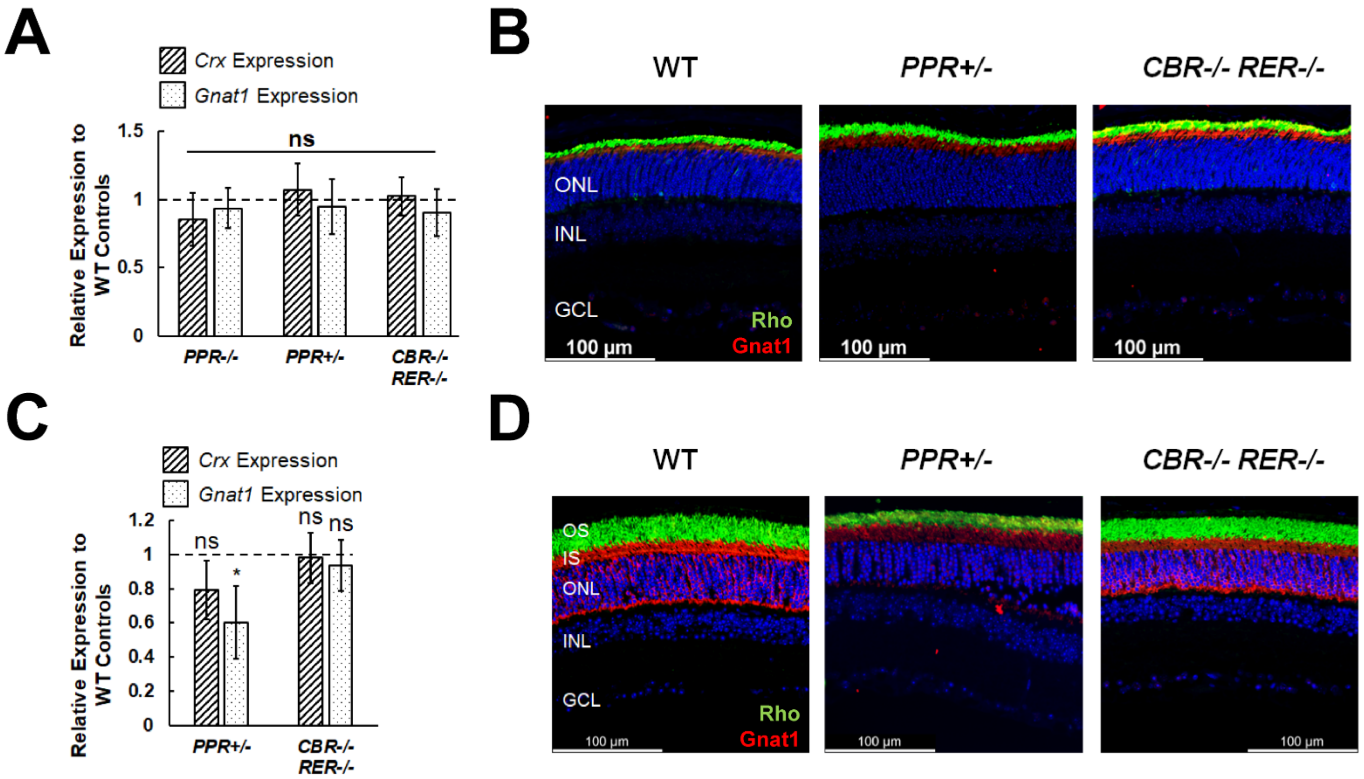


**Supplemental Figure 4. Expression of rod photoreceptor-related genes, *Crx* and *Gnat1* in mutant retinas. (A, C**) qRT-PCR analysis of *Crx* and *Gnat1* expression in retinal samples of the indicated mice at P14 (**A**) and 6MO (**C**). Results are plotted as relative expression to the *WT* controls. Statistics is done by one-way ANOVA with Tukey's multiple comparisons. ns means not significant. Asterisk (*) denotes $p \leq 0.05$. (**B, D**) Immunohistochemistry staining of Rho and Gnat1 on retinal cross-sections of the indicated mice at P14 (**B**) and 6MO (**D**). Scale bar represents 100 µm for all image panels.

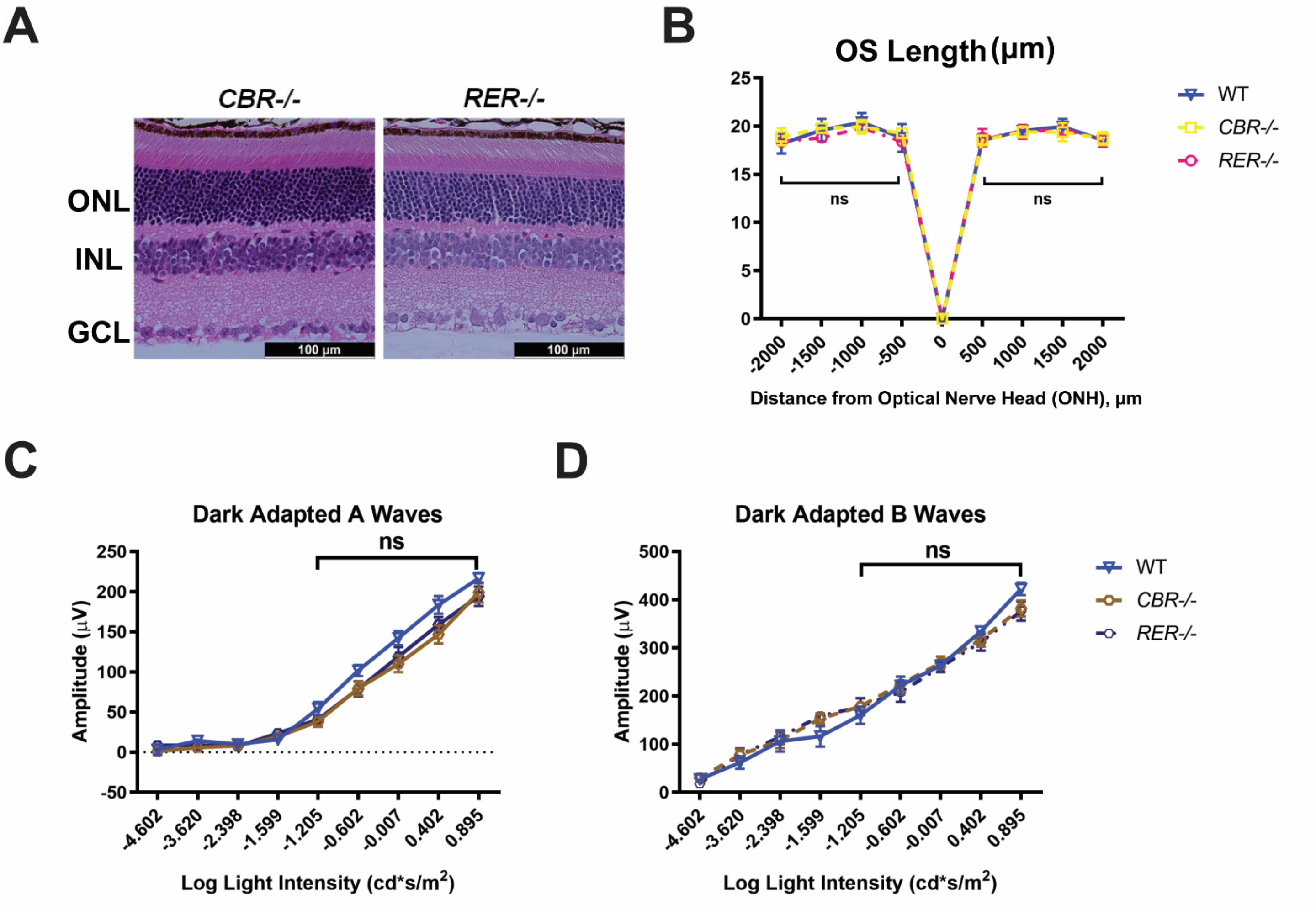


**Supplemental Figure 5. Knockout of individual enhancers does not impact retinal morphology and function.** (**A**) Hematoxylin and Eosin (H&E) cross-section staining of 6MO *CBR-/-*, and *RER-/-* retinas. ONL: outer nuclear layer; INL: inner nuclear layer; GCL: ganglion cell layer. Scale bar = 100 µm for all image panels. (**B**) OS thickness in 6MO *WT*, *CBR-/-*, and *RER-/-* retinas at various positions from the optic nerve head (ONH). Error bars represent mean (SD) (n ≥ 4). (**C, D**) Electroretinogram (ERG) analysis of 6MO *WT*, *CBR-/-*, and *RER-/-* mice, showing mean amplitudes (µV) of dark-adapted A-waves (**C**) and B-waves (**D**) at various stimulus light intensities. Error bars represent SEM (n ≥ 7). All statistics is done by comparing to *WT* control with two-way ANOVA with Tukey’s multiple comparisons. ns means not significant.

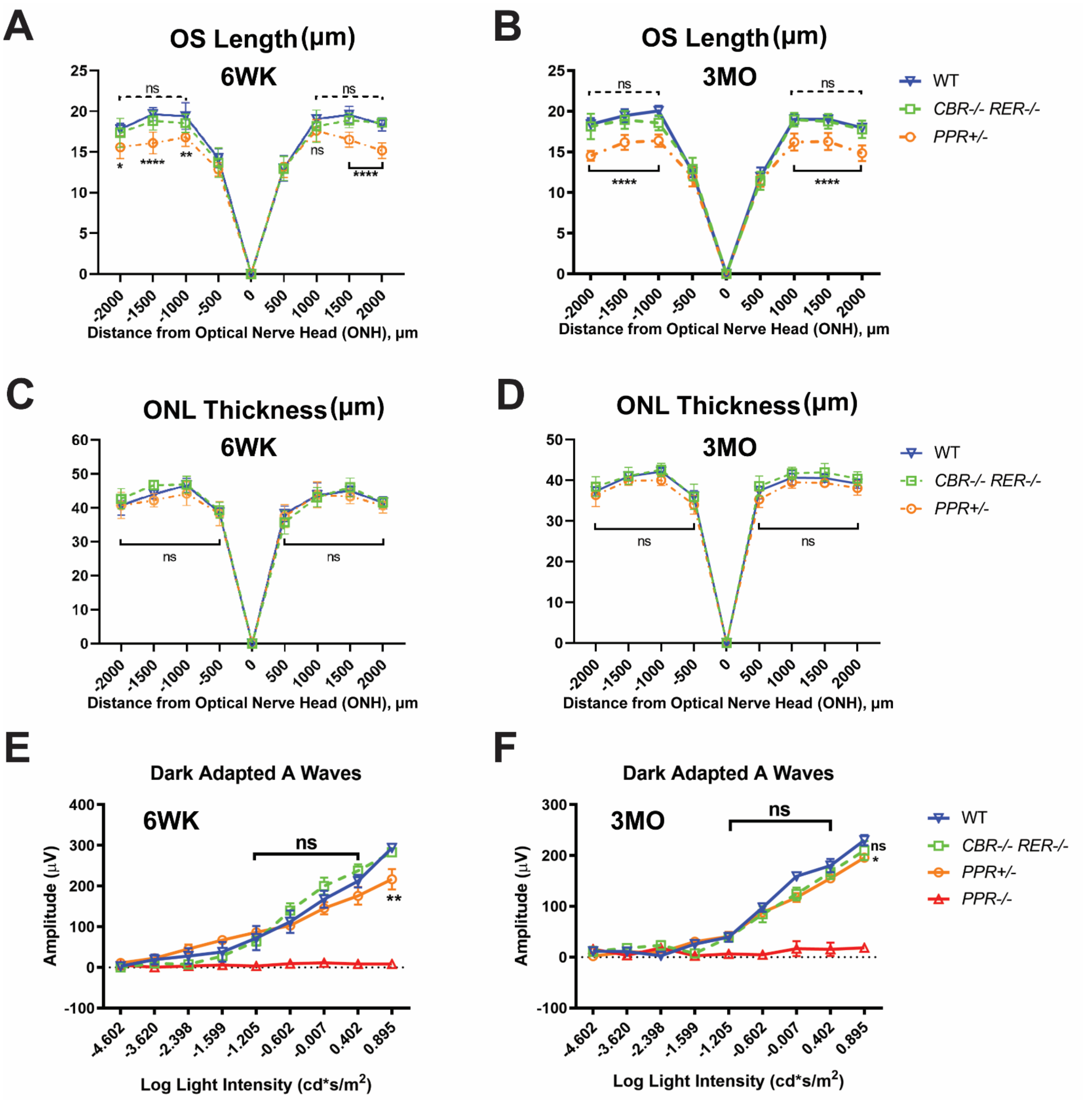


**Supplemental Figure 6**. **Knockout of both *Rho* enhancers (*CBR-/-RER-/-*) does not alter retinal morphology and function in young adults.** (**A** & **B**) OS thickness in 6WK (**A**) and 3MO (**B**) retinal samples of *WT* and mutants (n ≥ 4). Dashed black line represents the comparison between *WT* and *CBR-/-RER-/-* samples, and solid black line represents the comparison between *WT* and *PPR+/-* samples. (**C** & **D**) ONL thickness in 6WK (**C**) and 3MO (**D**) retinal samples of *WT* and mutants (n ≥ 4). Solid black line represents the comparison between *WT* and mutant samples. (**E** & **F**) Electroretinogram (ERG) analysis of dark-adapted A-waves for 6WK (**E**) and 3MO (**F**) samples of *WT* and mutants. Mean amplitudes (μV) are plotted against stimulus light intensity. Error bars represent SEM (n ≥ 7). All statistics is done by comparing to *WT* control with two-way ANOVA with Tukey's multiple comparisons. Asterisks (*, **, ****) denote $p \leq 0.05$, $p \leq 0.01$, $p \leq 0.0001$, respectively. ns means not significant.

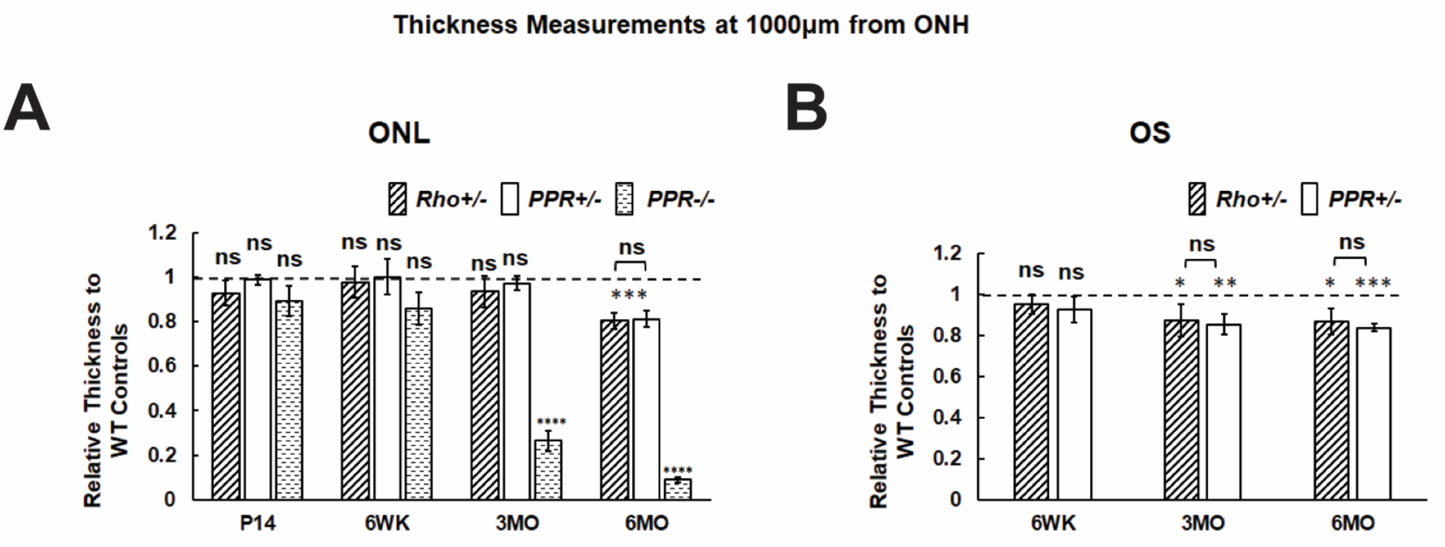


**Supplemental Figure 7. Comparable analysis of ONL and OS thickness in *Rho PPR* and *Exon* knockout mutants.** Bar graphs of ONL (**A**) and OS (**B**) thickness in *Rho+/-*, *PPR+/-*, *PPR-/-* retinas relative to *WT* control at different ages. Asterisks (*, **, ***, ****) denote p ≤ 0.05, p ≤ 0.01, p ≤ 0.001, p ≤ 0.0001, respectively by one-way ANOVA with Tukey's multiple comparisons. ns means not significant.
